# Proteomics reveals conserved features of histone biology in dinoflagellates

**DOI:** 10.64898/2026.10.29.761821

**Authors:** Nikolaj Brask, Ronie Haro, John M. Archibald, Courtney W. Stairs, Claudio H. Slamovits

## Abstract

Current knowledge of DNA packaging in eukaryotic nuclei largely centers on its basic unit, the nucleosome. Multiple lineages have, however, diverged from this paradigm, most notably dinoflagellates. Among other molecular oddities, these enigmatic protists have enormous genomes, permanently condensed chromosomes and a foreign gene encoding Dinoflagellate Viral Nucleoprotein (DVNP), which is used for packing DNA. Some dinoflagellates also have Histone-like Proteins (HLPs) acquired from bacteria via lateral gene transfer. Dinoflagellates were long thought to lack histones entirely, but canonical histone genes have been identified by transcriptomics. Here, we performed proteomics on purified nuclei and confirmed the presence of all four core histone classes, along with variants, and DVNP in the dinoflagellates *Oxyrrhis marina* and *Amphidinium carterae* and HLPs in *A. carterae*. We also identified putative histone post-translational modifications found in canonical eukaryotes. Comparative genomics of dinoflagellates revealed histone-handling and modification machinery alongside an apparent reduction in, or diversification of, machinery responsible for replication-dependent nucleosome deposition (CAF-1), consistent with a predominantly transcription-associated role for nucleosomes. Furthermore, proteomics revealed modifications of DVNP and HLP, raising the possibility of machinery co-opting. These observations support a new model of a dinoflagellate chromatin that includes the localization of canonical nucleosomes to “insulated chromatin territories” (INTs).

## INTRODUCTION

Few macromolecular structures are as faithfully conserved as the nucleosome. In nearly every eukaryote, DNA is wound around an octamer of the four core histones (H2A, H2B, H3 and H4) [1], proteins whose sequences are conserved across more than a billion years of evolution [2,3]. On top of these proteins sits a layer of regulation: the post-translational modification of histones, written, read and erased by enzymes that govern how the genome is folded, transcribed and repaired [4,5]. Because the system is nearly universal, its apparent abandonment in dinoflagellates is important for our understanding of chromatin structure, function and evolution. Dinoflagellates are aquatic protists that can cause harmful algal blooms, bioluminescence, and are the photosynthetic component of coral reefs [6,7]. They have some of the largest genomes known to biology, in some species exceeding the human genome many times over [8]. Their DNA is held in the dinokaryon, a nucleus unlike that of other eukaryotes: chromosomes remain condensed throughout the cell cycle in a birefringent, liquid-crystalline state, and histones are present at very low levels [9–11]. These histones were thought to have been displaced by two unrelated families of DNA-binding proteins: bacterial-type histone-like proteins (HLPs) and the Dinoflagellate Viral nucleoproteins (DVNPs), both acquired by lateral gene transfer (LGT). The acquisition of the genes for these proteins appears coincident with the loss of bulk nucleosomal packaging [10,12–14]. DVNP deviates from the viral counterpart by an intrinsically disordered N-terminal extension [15].

How large dinoflagellate genomes are packaged, once the nucleosome is set aside, is poorly understood. One prevailing model is that the chromosomes are organised as a cholesteric liquid crystal, with stacked layers of DNA fibrils rotating progressively from layer to the next, giving rise to the nested arches seen in electron micrographs [9,16,17]. Because the protein-to-DNA ratio in dinoflagellate chromosomes is roughly an order of magnitude lower than that of typical eukaryotic chromatin, condensation is thought to arise largely from the self-assembly of DNA at high concentration rather than from a protein scaffold [10,18]. While this model accounts for much of the observed ultrastructure, recent chromosome-conformation data have prompted a reappraisal of how strictly the liquid-crystal description holds [19–21].

Furthermore, histones were not eliminated in dinoflagellate, as once believed. Transcriptomic and genomic data reveal a full complement of core histone genes and histone modification enzymes [22,23], suggesting the lineage has not dispensed with the nucleosome. Evidence at the level of protein is scarce and fragmentary. Despite multiple attempts, immunodetection of core histones is largely confined to the nuclear periphery [24,25]. Mass spectrometry of the parasite *Hematodinium* sp. recovered peptides from histone H2A alone [10]; chromatin proteomics of *Lingulodinium polyedra* detected only histone H4, on the strength of two peptides [26]; and an enriched-nuclei survey of *Prorocentrum cordatum* detected H2A, H2B and H4 but not H3, each at low abundance [27]. Proteomics of an environmental bloom containing *Prorocentrum donghaiense* recovered all four classes but only H4 peptides were reported as unique to *Prorocentrum* [28]. Histone modifications have been largely detected with antibodies [24,29], and mass-spectrometric assignments rest largely on peptides conserved across eukaryotes [29]. All four core histones have yet to be detected from a dinoflagellate in culture under stringent criteria. The protein-level inventory of dinoflagellate chromatin is incomplete.

These ambiguities bear most heavily at the base of the dinoflagellate tree, where the evolutionary transitions to the dinokaryon occurred. Phylogenomic analyses consistently place the syndinians and the free-living marine heterotroph *Oxyrrhis marina* as early-branching lineages, diverging before the “core” dinoflagellates (Dinokaryota) in which the fully elaborated dinokaryon is found [30,31]. *Oxyrrhis* is therefore potentially highly informative: it is easily cultured and sits close to the root of the group and may preserve an ancestral, or at least intermediate, nuclear condition [32,33]. Recent volume electron microscopy reinforces this view, describing the *Oxyrrhis* nucleus as a midpoint between the typical eukaryotic nucleus and the dinokaryon; its chromosomes are condensed throughout the cell cycle, as in other dinoflagellates, yet are thinner and longer than those of core species and lack the characteristic arched banding pattern [34]. By contrast, *Amphidinium carterae* is a photosynthetic core dinoflagellate with a fully developed dinokaryon, for which transcriptome resources are available [35,36].

Here we use proteomics on whole cells of *Oxyrrhis marina*, and on isolated nuclei of *O. marina* and *A. carterae*, to characterize the histone proteins within dinoflagellates. Using expanded proteomics searches that include common variable modifications on histones, we identify all four core histone classes in both species, with candidate modifications on the recovered peptides. We also detect putative modifications on DVNP and HLPs. Using scanning and transmission electron microscopy (SEM and TEM), we assess the morphology of isolated *Oxyrrhis* nuclei and observe constricted bodies and pore-like surface features. The histone variants of *Oxyrrhis* show a high degree of conservation with reference histones from other eukaryotes. Among identified histone variants, a putative CENPA variant was found. Using phylogenetics, a histone H3 was found in most dinoflagellates sampled and branches with bona fide CENPA/CENH3 histones. Examination of gene density in two symbiodiniacean genome assemblies provides context for models in which histone-containing chromatin is concentrated near chromosome ends.

## RESULTS

### Proteomics reveals core histones in *O. marina* and *A. carterae*

Using data-independent and data-dependent acquisition (DIA and DDA), we generated proteomes of *O. marina* whole cells and nuclear preparations, respectively, identifying 7,558 proteins in the whole-cell sample and 1,985 proteins in the three nuclear preparations (R1-3). For both samples, most identified proteins could be functionally annotated using InterProScan [37] with a threshold of E ≤ 10^−5^ (5,175 of 7,558 proteins (68.5%) in the whole-cell sample and 1,592 of 1,985, 80.2%, in R1). Functionally annotated proteins also contributed most of the measured protein intensity (69.6% and 88.9%, respectively) [Figure 1B; Table S1]. These results indicate that much of the experimentally accessible proteome has recognisable sequence features. Proteins predicted by DeepLoc 2.0 [38] to be nuclear contributed 9.3% of summed intensity in R1, compared with 5.5% in the whole-cell sample, a 1.7-fold increase [Figure 1B]. Proteins predicted to be cytoplasmic nevertheless contributed 57.3% of R1 intensity, consistent with substantial carry-over of non-nuclear material.

**Figure 1.**
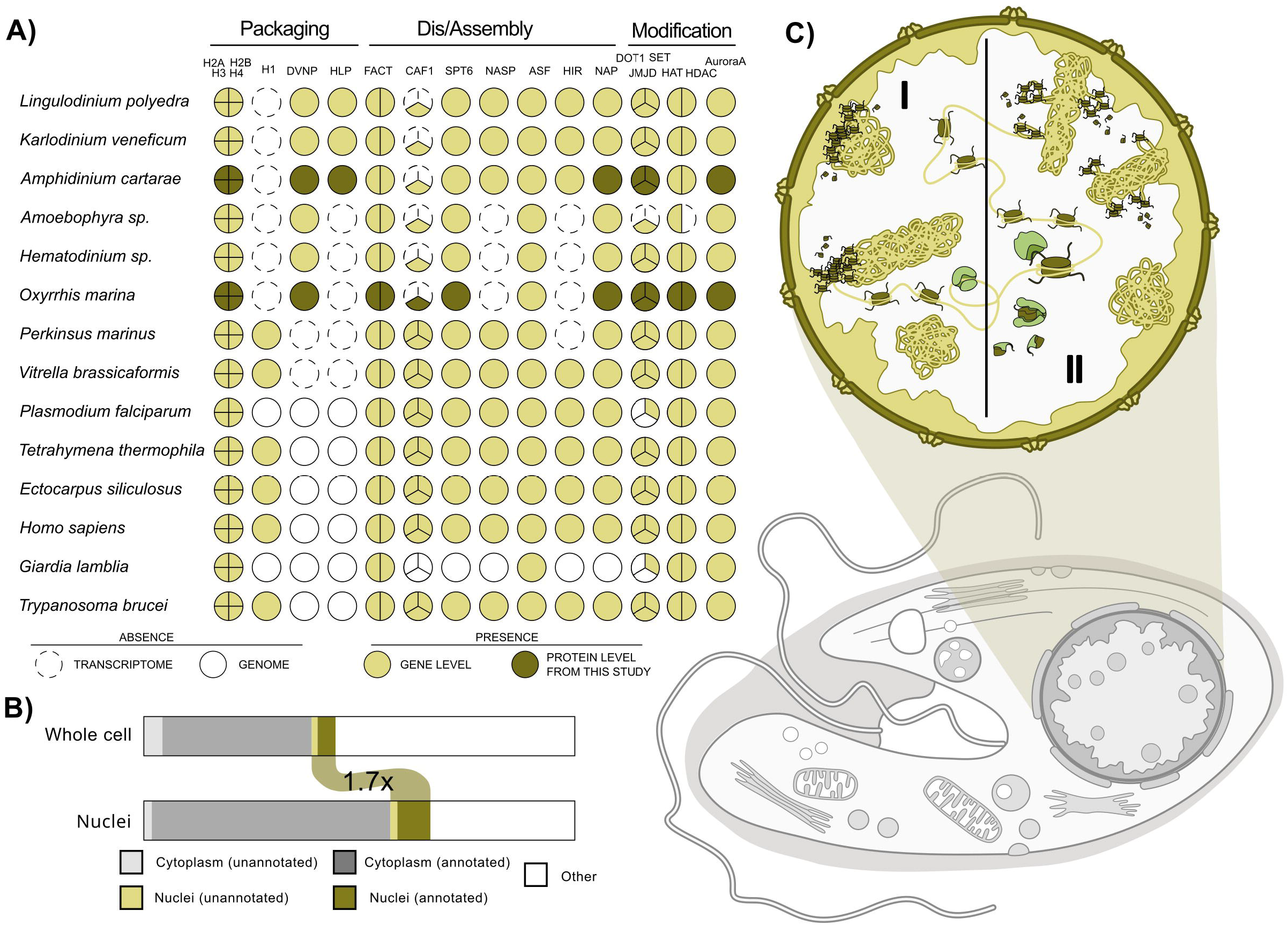
Nucleosome, DNA-packaging and chromatin-machinery components in dinoflagellates and other eukaryotes, and proteome composition of *O. marina* whole cells and nuclei. (A) Coulson plot showing the presence or absence of core histones (H2A, H2B, H3 and H4), linker histone H1, DVNP and HLP (packaging), histone chaperones and assembly factors (FACT, comprising SPT16 and SSRP1; CAF-1, comprising FAS1/Cac1, FAS2/Cac2 and FAS3/RBBP4; SPT6; NASP; ASF1; HIR/HIRA; and NAP) (dis/assembly), and histone-modifying enzymes (DOT1, SET-domain and JmjC-domain proteins, HATs, HDACs and Aurora A kinase) (modification) in six dinoflagellates and eight other eukaryotes. Segments of divided circles represent individual components. Light fill, present at the gene level; dark fill, also detected at the protein level in this study; open circles, not detected in a genome; dashed circles, not detected in a transcriptome. Accessions and sequences are given in Table S4. (B) Proportion of summed razor intensity in the *O. marina* whole-cell sample and the most histone-enriched nuclear preparation (R1) contributed by proteins predicted by DeepLoc to localise to the cytoplasm or nucleus, each subdivided into proteins with and without InterProScan annotation; “Other” comprises all remaining predicted localisations. Proteins predicted to be nuclear contributed 1.7-fold more of the summed intensity in R1 than in the whole-cell sample. (C) Proposed “insulated nucleosome territories” model: bulk DNA is packaged by DVNP/HLP and nucleosomes persist is only (I) at chromosome ends anchored at the nuclear envelope [24] and (II) over extrachromosomal loops, with machinery and chaperones (green) detected in this study

Within the proteomic searches of both dinoflagellate species, all four core histones could be identified. In *O. marina*, the relative contribution of core histones to total quantified protein intensity increased in nuclear preparations. Using razor-peptide IonQuant intensities from the initial searches, summed histone intensity accounted for approximately 0.033% of whole-cell intensity and 2.26%, 0.29% and 0.079% in nuclear preparations R1, R2 and R3, respectively. Relative to the whole-cell sample, these correspond to approximately 70-, 8.9- and 2.4-fold enrichment [Figure 2A; Table S2], or 49-, 5.8- and 1.6-fold using the expanded histone searches. In the *A. carterae* nuclear preparation, core histones accounted for 0.094% of summed intensity [Table S3]; no whole-cell *A. carterae* sample was available for comparison.

**Figure 2.**
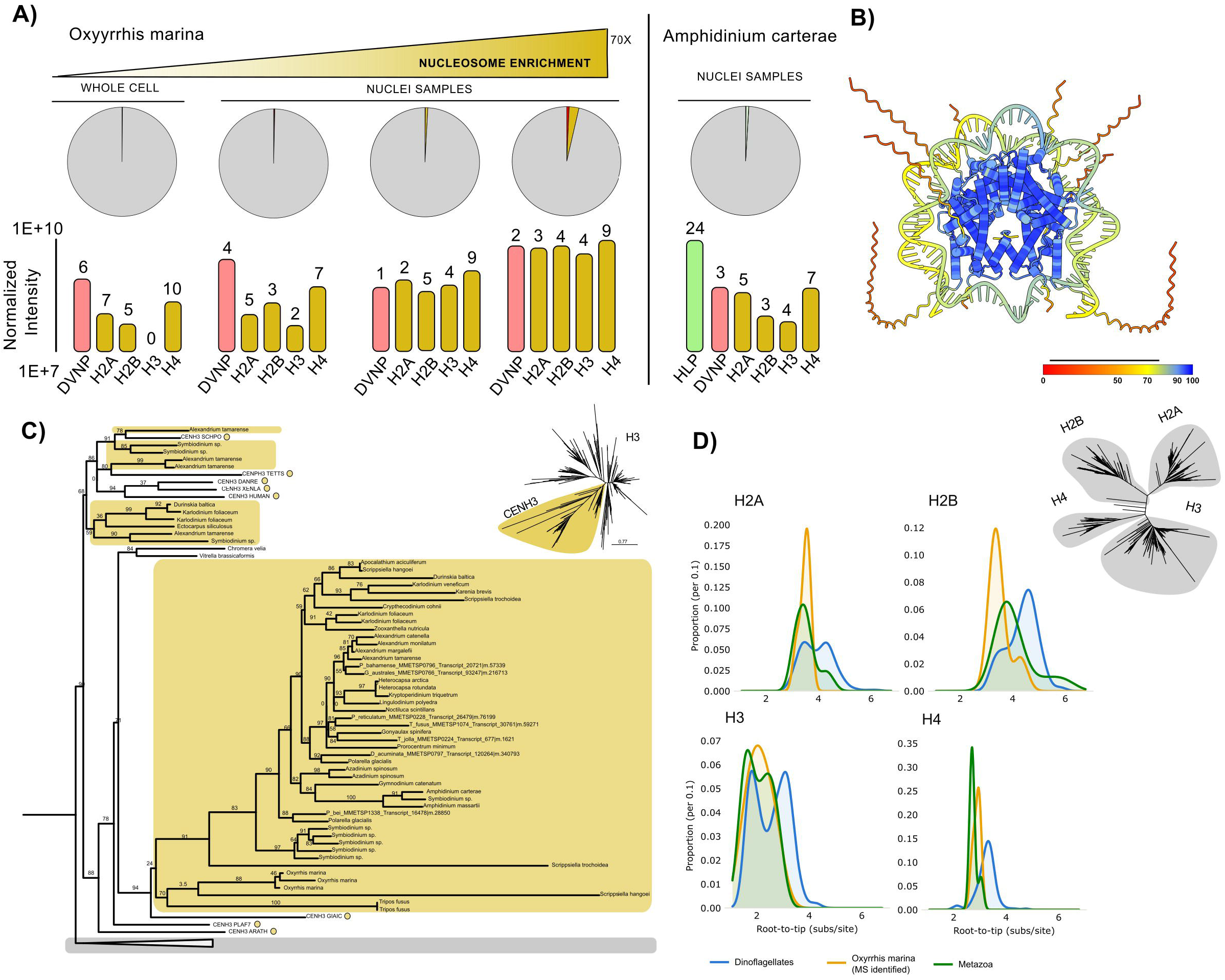
Core histones and DVNP in the *O. marina* and *A. carterae* proteomes, and histone H3 phylogeny. (A) Pie charts show the proportion of summed razor intensity attributed to core histones (yellow) and DVNP (red) in the *O. marina* whole-cell sample and in nuclear preparations ordered by increasing histone enrichment (R3, R2 and R1; up to approximately 70-fold relative to whole cells), and in the *A. carterae* nuclear preparation. Bar charts show intensities of DVNP, HLP (green) and each core histone class, normalised so that each sample has the same summed intensity as the whole-cell sample (log scale); numbers above bars give razor peptides in the initial searches. (B) Predicted structure of a histone octamer with DNA, coloured by per-residue confidence (0–100). (C) Maximum-likelihood phylogeny of histone H3 sequences from dinoflagellates and other eukaryotes. Shaded boxes indicate dinoflagellate-containing clades, circles mark reference centromeric H3 (CenH3) sequences and numbers at nodes give branch support. Inset: unrooted tree with the CenH3-containing clade shaded. (D) Root-to-tip distances per histone family in a maximum-likelihood tree of all four families (inset) for dinoflagellates (blue), proteomics-identified *O. marina* histones (orange) and metazoa (green).

Using FragPipe searches against the predicted proteome databases of both sampled dinoflagellates, we identified all four core histone classes, H2A, H2B, H3 and H4. At least one predicted sequence from each class was supported by two or more peptides in at least one sample [Figure 2A; Tables S2, S3]. In both species, each core histone class was represented by at least one detected peptide that is not identical to any human sequence [Figures S2, S3]. *O. marina* H2A and H2B sequences are divergent from human histones, whereas conserved regions of H3 and H4 can yield peptides shared with human homologues.

In the most histone-enriched *O. marina* preparation (R1), the summed intensity ratio was approximately 1:1.1:0.71:1.6 for H2A, H2B, H3 and H4, respectively. These similar signal magnitudes are compatible with recovery of all four histone classes, although do not precisely measure nucleosome stoichiometry. Across the *O. marina* samples, three predicted H2A proteins, three H2B proteins, two H3 proteins and three H4 proteins were identified.

For representative *O. marina* sequences in R1 (expanded histone search), H2A (MMETSP0469 g.18454) had three reported peptides and 17.83% coverage, H2B (MMETSP1426 g.61156) three peptides and 21.55% coverage, H3 (MMETSP0469 g.127961) six peptides and 37.31% coverage, and H4 (MMETSP0469 g.88073) eight peptides and 37.74% coverage. In *A. carterae*, representative H2A (MMETSP0259 g.2185), H2B (MMETSP0259 g.83761), H3 (MMETSP0259g.72084) and H4 (MMETSP0259 g.31896) sequences had four, four, five and eleven reported peptides, with 20.44%, 18.49%, 26.49% and 56.20% coverage, respectively [Tables S2, S3]. Coverage therefore varied substantially among histone classes and predicted sequences.

### Nucleosome assembly and modification machinery widespread in dinoflagellates

A bioinformatic survey revealed that in addition to core histones, dinoflagellates retain components of the machinery for nucleosome assembly and modification [Figure 1A; Table S4]. Homologues of the two FACT subunits, SPT16 and SSRP1, and the histone chaperone NAP were recovered across the dinoflagellate taxa surveyed. Other chaperones have a more restricted distribution: HIR/HIRA homologues were recovered from several core dinoflagellates, including *A. carterae*, but were missing in the *O. marina* transcriptome analysed, whereas ASF1 was found in all six. In *O. marina*, SPT16, SSRP1, SPT6 and NAP were also detected at the protein level [Figure 1A].

Using homology and HMM-based searches, homologues of the two larger chromatin assembly factor-1 (CAF-1) subunits, FAS1/Cac1 and FAS2/Cac2, could not be identified from the dinoflagellate datasets surveyed. The small subunit MSI1/RBBP4 was recovered from several taxa, including *O. marina*, in which RBBP4 was also detected at the protein level. This pattern is broadly consistent with previous comparative analyses [22,23]. We also recovered candidate histone-modifying enzymes, including HDAC and DOT1 homologues in both focal species; DOT1 was detected at the protein level in both, and HDAC in *O. marina* [Figure 1A]. These searches establish a candidate repertoire; while non-detection in a transcriptome does not by itself establish gene loss, BUSCO completeness of the genome assemblies for both genera was 86–100% [39,40; Figure S1], suggesting that the undetected genes are indeed not present.

### Histone conservation

The detected histones contained conserved core regions. In *O. marina*, the H3 sequence MMETSP0469 g.127961 retains a recognisable canonical-like N-terminal tail and histone-fold region and differs from human H3.1 at only four positions (with a −1 offset), whereas other predicted histones contain altered or extended termini. Similarly, the most frequently detected *O. marina* H4 sequence (MMETSP0469 g.88073) differs from human H4 at only two positions (with a 3-residue N-terminal extension), so most of its tryptic peptides are shared with human H4. *A. carterae* histones likewise retain recognizable core regions despite terminal divergence. Conservation of these regions is consistent with retention of histone interactions, but the sequences alone do not establish assembly into an octamer or the function of individual variants. A structure prediction of a histone octamer assembled from proteomic *O. marina* histones with DNA places the histone-fold domains with high per-residue confidence and the disordered terminal tails with low confidence [Figure 2B]. In a phylogeny of H3 sequences [Figure 2C], many dinoflagellates, including *Oxyrrhis marina* and *Amphidinium carterae*, group within the clade containing all validated centromeric H3 (CenH3) sequences from representative organisms [2,3]: fission yeast, *Tetrahymena*, vertebrates, *Plasmodium* [41] and *Giardia* [42]. This suggests the presence of CenH3 variants in dinoflagellates. Whilst most dinoflagellate histones have long root-to-tip distances in the histone phylogenetic tree, proteomic-identified *O. marina* histones generally have shorter root-to-tip distances than other dinoflagellate histones [Figure 2D].

### Proteomics reveals canonical histone post-translational modifications in dinoflagellates

Using an extended search of common histone variable modifications and increased missed cleavages, we recovered candidate methylated, dimethylated, acetylated and phosphorylated histone peptides in *O. marina* and *A. carterae* [Figure 3; Tables S2, S3]. These assignments describe peptide-spectrum matches (PSMs) consistent with modified peptides.

**Figure 3.**
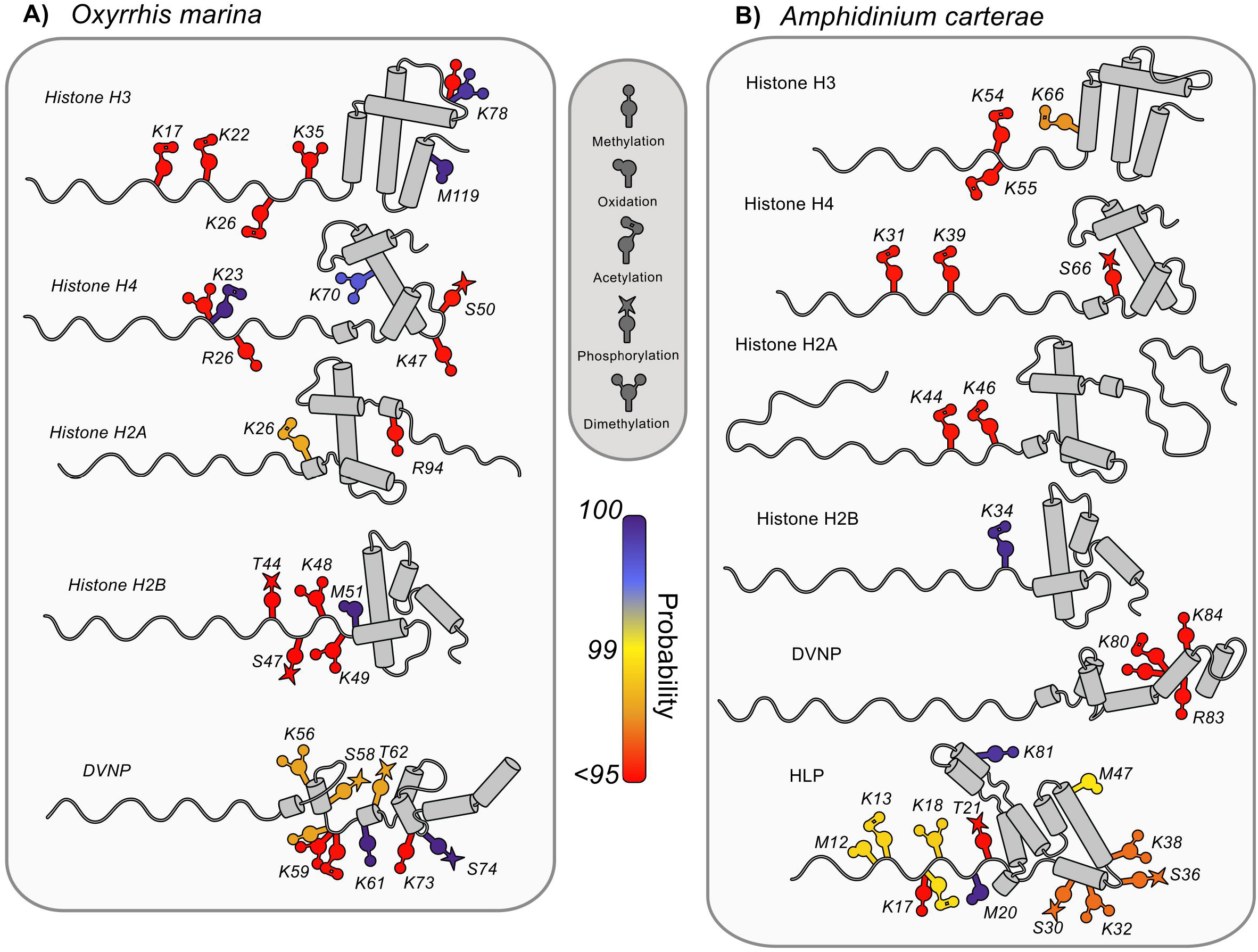
Candidate post-translational modifications on dinoflagellate histones, DVNP and HLP. Candidate PTMs identified in expanded FragPipe searches are shown on cartoon representations of representative histone, DVNP and HLP sequences from *O. marina* (left) and *A. carterae* (right); cylinders indicate α-helices. Symbols denote methylation, oxidation, acetylation, phosphorylation and dimethylation, and colour indicates the probability of the highest-scoring PSM carrying each modification (<95% to 100%). Residue numbers refer to the predicted sequences in Tables S2 and S3.

Histone H3 of *O. marina* had candidate dimethylation at K78 of the predicted sequence MMETSP0469 g.127961, which corresponds to the human H3 residue K79. This assignment was reported in two nuclear preparations, R1 and R3, and in the whole-cell sample, and monomethylation of the same residue was assigned in R1 [Figure 3]. The modified peptide, EIAQDFKTDLR, is identical to its human counterpart [Figure S2]. The same sequence had candidate acetylation at K17 and K22 (conventional H3K18 and H3K23) in R1 and the whole-cell sample, and candidate acetylation at K26 (H3K27) and dimethylation at K35 (H3K36) in R1, all with lower PSM probabilities (<95%) [Figure 3]. In *A. carterae*, candidate acetylation was assigned at K54, K55 and K66 of H3 (K54 and K66 align with H3K14 and H3K27), and methylation at K93 of a second H3 sequence.

Histone H4 of both dinoflagellates had candidate modifications in the N-terminal region and structured domain. In the *O. marina* whole-cell sample, phosphorylation was assigned at S65 of MMETSP1424 m.58436, equivalent to S50 of the near-identical sequence MMETSP0468 g.21715 shown in Figure 3, and *A. carterae* H4 carries a candidate S66 phosphorylation assignment. Pairwise alignment places both sites at the position of human H4S47. At the position of H4K20, *O. marina* g.21715 K23 was assigned dimethylation in R1, and the equivalent residue of m.58436 (K38) acetylation in the whole-cell sample, on a peptide that is not shared with human H4 [Figure S2]; methylation was also assigned at the adjacent R26 (H4R23) in R1. *A. carterae* H4 had candidate acetylation at K31 and K39, which align with H4K12 and H4K20, respectively. Additional candidate methyl modifications in the *O. marina* H4 histone-fold domain included K70, a position occupied by arginine (R67) in human H4, supported by a species-specific peptide [Figures 3, S2].

### DVNP and HLP abundance and modifications in dinoflagellate nuclei

Alongside the core histones, we identified DVNP in the *O. marina* nuclear and whole-cell proteomes and in the *A. carterae* nuclear proteome. In the most histone-enriched *O. marina* sample (R1), the DVNP-to-H2A-to-H2B-to-H3-to-H4 intensity ratio was approximately 1:0.9:1.0:0.6:1.4. DVNP was supported by two razor peptides in R1, one in R2, four in R3 and six in the whole-cell sample in the initial searches [Figure 2A]. Its normalised intensity was approximately 7.6-fold higher in R1 than in the whole-cell sample, compared with approximately 70-fold enrichment of summed histone intensity. In *A. carterae*, HLPs dominated the detected packaging-protein signal: seven HLP sequences, supported by 24 razor peptides, accounted for 0.81% of summed intensity, compared with 0.094% for core histones and 0.049% for DVNP [Figure 2A; Table S3].

Candidate DVNP modifications included acetylation and methylation in both species, with dimethylation and phosphorylation additionally assigned in *O. marina*, almost entirely in the whole-cell sample [Figures 3, S4]. Of the seven HLP sequences detected in *A. carterae*, three carried candidate acetylation, methylation, dimethylation and phosphorylation assignments, and a fourth carried methylation [Figure 3; Table S3].

### Electron microscopy of isolated nuclei show unique features

Using electron microscopy, we examined the morphology of *O. marina* nuclei isolated as for proteomics but with an additional Percoll/sucrose density-gradient purification step [Figure S5]. In the TEM images, membrane boundaries were poorly resolved, although numerous chromosomes appeared to congregate around the nuclear periphery [Figure S5]. Scanning electron microscopy revealed a heterogeneous field of discrete, roughly spheroidal to ovoid bodies [Figures S5, S6]. Two recurring surface morphologies were observed: relatively smooth or folded surfaces, and rougher, granular surfaces with variable lobing [Figure S6A, B].

Several bodies exhibited paired profiles or circumferential constrictions [Figures S5, S6A, B]. These could represent division-related morphologies, although apposition, fragmentation and preparation-induced deformation cannot be excluded. Some smoother surfaces contained pore-like depressions or openings (arrows in Figure S5; Figure S6C). Homologues of nucleoporins from the outer-ring, inner-ring, central-channel and cytoplasmic-filament subcomplexes of the nuclear pore complex, including Nup84, Nup96, Nup120, Seh1, Sec13, Nic96, Nup155/170, Nup62, Nup54/57 and Dbp5, were detected in the *O. marina* proteome [Table S5], consistent with the presence of pore complexes in the nuclear material.

### Gene density increases towards contig ends in symbiodiniacean genomes

Because chromosome-end chromatin has been proposed as a principal location of dinoflagellate nucleosomes [24], we examined how genes are distributed along contigs in two highly contiguous dinoflagellate genome assemblies, those of *Cladocopium goreaui* and *Durusdinium trenchii*. In both species, gene density was elevated towards contig ends [Figure S7]. In *C. goreaui*, smoothed density rose from approximately 36–37 genes per Mb in contig interiors to approximately 39 genes per Mb near the ends, and in *D. trenchii* from approximately 31 to 35–37 genes per Mb (Kolmogorov–Smirnov D = 0.014, *p* = 6.6 × 10^−5^, and D = 0.020, *p* = 7.1 × 10^−19^; paired Wilcoxon *p* = 7.9 × 10^−3^ and 3.3 × 10^−17^, respectively). Contig ends do not necessarily correspond to chromosome ends, so these patterns indicate gene-rich terminal regions only where assemblies approach chromosome scale.

## DISCUSSION

### Nucleosome evolution within dinoflagellates

The lysine-rich, low-complexity tails of most of histone H1 proteins evolve rapidly and lack the conserved residues needed for reliable homology detection [43]. Previous studies have proposed a loss of histone H1 among dinoflagellates [44,45], although Marinov and Lynch reported putative linker histones from many dinoflagellate transcriptomes and attributed failures to detect them to incomplete genome assemblies [23]. No confident H1 homologue was recovered from any dinoflagellate in our survey [Figure 1A]. Loss of histone H1 is not unprecedented, although it is mostly known from unrelated parasitic lineages such as *Giardia* and *Plasmodium*. Neither lineage encodes a recognisable H1 [46,47], and in *Plasmodium* chromatin regulation relies heavily on histone modifications within a largely euchromatic genome [47]. In trypanosomes, histone H1 lacks the globular domain seen in most eukaryotes and is composed mostly of lysine, alanine, valine and proline [43,48]. These lineages show that H1-mediated higher-order packing is not a eukaryotic necessity, and that its absence does not compromise the composition of the nucleosome-interacting machinery.

Alongside H1, CAF-1 is also widely absent from dinoflagellate genomes and transcriptomes. Both HMMs and BLAST-based homologue searching fail to find two of the three subunits, returning hits for only RBBP4. RBBP4 is not specific to CAF-1 and can participate in other chromatin-associated complexes [49], so its retention alone, even with protein-level detection in *O. marina*, does not establish an intact CAF-1 complex. Failure to recover the two larger CAF-1 subunits is consistent with reduced reliance on conventional replication-coupled histone deposition [23]. However, transcriptome incompleteness, sequence divergence or use of alternative chaperones could account for absences seen. The same qualification applies to the non-detection of HIR/HIRA in *O. marina*. FACT and NAP homologues, both detected at the protein level in *O. marina*, support the retention of histone-handling capacity, while HIR/HIRA in several core dinoflagellates provides a candidate route for replication-independent deposition. These factors are not exclusive markers of transcription: histone exchange and chromatin restoration can also accompany DNA repair and other DNA transactions [50,51]. Dinoflagellates are the only group of the surveyed organisms that have partial absences in the CAF-1 complex, and previous studies have shown conserved presence of CAF-1 across multiple eukaryotic supergroups including alveolates (*Plasmodium*), although no dinoflagellate was included in the analysis [52]. In our survey, complete CAF1 absence is seen in the diplomonad *Giardia*. Searches of multiple InterPro/Pfam entries [53] identified no CAF-1 component in any diplomonad. Although CAF1 is required for many organisms’ development and gene expression, there is some degree of redundancy between HIR and CAF-1, creating plasticity in nucleosome deposition [54–57].

### Dark genes and the proteome–transcriptome discrepancy

Dinoflagellate transcriptomes contain many predicted proteins without recognised functions, including a substantial complement of conserved dark (unannotated) genes in dinoflagellates [58,59]. The greater apparent annotatability of the detected proteome may reflect preferential recovery of abundant, conserved proteins with recognisable domains. It may also reflect differences in annotation criteria, transcript fragmentation and the inclusion of low-abundance or condition-specific transcripts in sequence assemblies. Thus, the proteome–transcriptome discrepancy cannot alone establish that unannotated genes are weakly expressed or dispensable. A matched analysis using the same reference database and annotation criteria would help distinguish biological differences from detection and annotation biases.

### Electron microscopy and proteome of isolated nuclei detect nuclear pores

The heterogeneous morphology of the nuclear preparations, even after the additional density-gradient purification used for electron microscopy, limits interpretation of the proteomic fractions as purified nuclei. The second population of electron-dense or granular structures [Figure S6] could include altered nuclei, other subcellular material or preparation artefacts. Enrichment of histone signal and the 1.7-fold increase in signal from proteins predicted to be nuclear are consistent with the intended isolation, but do not establish purity or the intranuclear location of each protein. Similarly, pore-like openings and constricted bodies should remain provisional morphological observations until supported by correlative imaging or molecular labelling, although detection of nucleoporins in the proteome [Table S5] supports the presence of pore complexes in the preparations. These preparations provide a route to recovering nuclear proteins, while preserving a distinction between biochemical enrichment and localisation within intact cells.

### Conserved histone features and variants

Our phylogeny places several dinoflagellate H3 sequences, including those of *O. marina* and *A. carterae*, within the clade containing validated CenH3 sequences [Figure 2C]. Because both CenH3 and the divergent, often extended dinoflagellate H3 variants evolve rapidly, this placement could be artifactual (e.g., suffering from long-branch attraction [60]). Proteomic searches of *O. marina* recovered a single peptide from one candidate CenH3 (MMETSP1424 m.15197, R3; Table S2), below our two-peptide threshold for a confident identification. Independent of tree topology, this sequence has CenH3 features: a longer N-terminal extension and loop 1 than that of canonical H3. In CENP-A, loop 1 forms part of the CENP-A targeting domain (CATD) required for centromere targeting [61]. If confirmed, a CenH3 would imply a dedicated centromeric nucleosome population in dinoflagellates, and hence a role for nucleosomes in chromosome segregation as well as transcription. This would fit the observation that dinoflagellate chromosome ends interact with extranuclear mitotic microtubules at the nuclear envelope [24,62], an anchorage at which centromeric nucleosomes could reside.

### Significance of histone PTMs

The candidate H3K79 dimethylation is of particular interest because it occurs in a conserved histone-core region. Its repeated assignment in *O. marina*, together with a DOT1 homologue detected at the protein level [Figure 1A], is consistent with retention of an enzymatic histone-modification pathway [63,64]. H3K79 and H3K36 methylation are associated with transcription elongation [65–67] and both residues are retained in most dinoflagellate H3 variants [23] (methylation at both has also been detected with antibodies in *A. pacificum* [29]); the candidate H3K79me2 (K78 in MMETSP0469 g.127961 and lower-probability H3K36me2 (K35) assignments in *O. marina* therefore fit a transcription-associated role for nucleosomes. Structural work on DOT1 provides a mechanistic precedent for recognition of a nucleosomal substrate [68]. Species attribution of the modified peptide remains particularly important for this conserved histone, because the modified peptide, EIAQDFKTDLR, is identical to its human counterpart and contamination cannot be excluded [Figure S2]. Nevertheless, an antibody-based detection of H3K79 methylation in *A. pacificum* [29], supports a dinoflagellate assignment to this peptide. The candidate phosphorylation in the conserved H4S47 region provides a more specific mechanistic connection to replacement histone deposition. In human cells, PAK2-mediated H4S47 phosphorylation promotes HIRA-mediated H3.3–H4 assembly and reduces CAF-1-associated H3.1–H4 deposition [69]. A modification at the equivalent dinoflagellate residue would therefore be compatible with the proposed bias towards replication-independent histone handling. This is a mechanistic analogy, not evidence that the complete pathway is conserved.

Candidate modifications at the position of H4K20 in both species, including on a peptide unique to *O. marina* H4, are of particular interest because H4K20 methylation states are coupled to DNA replication, repair and chromatin compaction in other eukaryotes [70]. PSM probabilities do not directly measure PTM localisation confidence, and searches allowing several modifications per peptide increase the number of possible assignments. The current dataset therefore supports prioritising specific sites for validation rather than inferring a complete regulatory programme. Confirmed modification of species-specific peptides, followed by mapping of histone occupancy relative to transcription, would connect the biochemical observations to the proposed role.

### DVNP modification and domestication

The lower apparent enrichment of DVNP relative to histones in some preparations (7.6-versus approximately 70-fold in R1) may reflect differences in extraction, peptide recovery and quantification. Lysine-rich sequences can yield short tryptic peptides, and extensive modification can distribute signal among several peptide forms. DDA and DIA also differ in sampling and quantification, making their raw intensity estimates unsuitable for direct comparison without harmonised processing. The additional HLP repertoire in *A. carterae* further complicates comparison with *O. marina*. Accordingly, similar DVNP and histone signals in these datasets need not contradict a major role for non-histone proteins in bulk chromosome organisation, especially as the basic chromosomal protein of *O. marina*, Np23 (DVNP), was isolated from nuclei as a dominant species [71].

Phosphorylation of DVNP was previously reported in *Hematodinium* [10], providing a precedent for modification of this protein family. The additional candidate PTMs reported here raise the possibility that DVNP function is regulated post-translationally. Experiments expressing DVNP in yeast showed nucleosome displacement and disruption of chromatin function [15]; whether particular DVNP modifications modulate DNA binding, and abate deleterious effects to expression, remains unknown. Such modulation could facilitate local access for transcription, replication or repair without requiring wholesale chromosome decondensation. The occurrence of similar modification classes on histones, DVNP and HLP is compatible with shared modifying enzymes, and could be a result of enzyme co-option.

The repeat-rich, low-complexity N-terminal region of *O. marina* DVNP could provide a substrate for regulatory diversification. Repeat expansion provide a possible route to generating additional modifiable residues, while acquisition of a histone-derived segment is another. The N-terminal region of DVNP is rich in basic residues and, in the *O. marina* sequences, contains tandem (ATRK)n repeats. Such repeats could provide multiple recognition sites for modifying enzymes without each site having to evolve independently. Alternatively, acquisition of a histone-derived N-terminal segment would provide pre-existing modification sites. An alignment of *O. marina* DVNP with viral DVNPs [Figure S8] shows that the detected peptides cover part of the conserved domain, and several modified residues are conserved in the viral sequences (e.g., T62 in all six and K61 in four of the six viral DVNPs aligned). This raises the possibility that modification of these sites predates the acquisition of DVNP by dinoflagellates. Viral proteins are commonly modified by host enzymes [72], and some large DNA viruses of algae encode their own chromatin-modifying enzymes, such as the SET-domain histone methyltransferase of Paramecium bursaria chlorella virus 1 [73]. Characterising DVNP modification in a DVNP-encoding virus and its host could resolve whether modification is ancestral or arose during domestication.

### Insulated nucleosome territory model for the dinoflagellate nucleus and its implications

A spatially restricted histone system is supported by the chromosome-end study of Kwok et al. [24]. Their anti-H3K9me3 immunofluorescence in *Karenia brevis* showed prominent peripheral labelling with little apparent signal over the central condensed chromosomes (their Figure 4A), providing spatial evidence that at least this modified H3 population is unevenly distributed within the nucleus. In *Crypthecodinium cohnii*, telomeric-repeat hybridisation to MNase-resistant material, together with anti-H2A co-immunoprecipitation of H2B and H3, provided complementary evidence for histone-containing telomeric chromatin [24]. These findings help reconcile low histone abundance with local nucleosome retention alongside extensive non-histone chromosome condensation. However, H3K9me3 immunofluorescence maps a modified histone population, not total nucleosome occupancy or active transcription, and the authors noted possible displacement of the nuclear envelope during fixation. Our analysis of two symbiodiniacean genome assemblies is relevant here: gene density increases towards contig ends [Figure S7], so terminal chromatin may be gene rich. If contig ends approximate chromosome ends, nucleosomes concentrated near chromosome termini could coincide with gene-dense, transcribed regions, partly reconciling the chromosome-end and transcription-associated models. The transcription-focused model proposed here remains a testable interpretation of the retained histone-handling machinery and candidate PTMs, alongside evidence for chromosome-end functions.

Recovery of all four core histones, CENPA candidates, histone PTMs and a retained subset of histone chaperones and modifying enzymes leads us to propose the “insulated nucleosome territory” (INT), model of dinoflagellate chromosomes [Figure 1C]. In this model, bulk chromosomal DNA is condensed without nucleosomes by DVNP and, in core lineages, HLPs [10,15,27], consistent with the low histone abundance we measured and with the dominance of HLPs in the *A. carterae* nuclear proteome. Nucleosomes persist in INTs, and function in a transcriptional role. The machinery (FACT, SPT6 and NAP) we detected as proteins act on histones providing a route to nucleosome assembly and regulation of expression. H3K79 methylation, catalysed by DOT1 (also detected) and found in *A. pacificum* [29], could mark elongating chromatin, consistent with the proposal that transcription proceeds through nucleosomes in dinoflagellates [23]. At chromosome ends anchored at the nuclear envelope, telomeric nucleosomes have been described [24]. This is one of two tentative locations for INTs; the modest gene enrichment near contig ends [Figure S7] suggests that gene-rich chromosome termini at the nuclear envelope could carry much of the nucleosomal chromatin. These must be re-established around each cycle of DNA replication but can be mediated by CAF1 alternatives (HIRA). INTs near the nuclear envelope could address the continued role of CENPA during closed mitosis, as the nucleosomes are now positioned where chromosomes are accessible to the cytoplasmic kinetochore in core dinoflagellates [62]. The available evidence suggests that histone supply is at least partly uncoupled from replication. Histone expression was stable across the cell cycle in cultured *A. pacificum* [25], whereas the S-phase rise and G2/M peak of histone signal reported from a *P. donghaiense* bloom [28] cannot be attributed to that species with confidence. The near-canonical histones of *O. marina* [Figure 2D] further suggest that the nucleosome core remained under strong constraint after DVNP was acquired, with greater divergence in the histones of most other dinoflagellates. The model predicts that INTs are concentrated over extrachromosomal loops and at nuclear envelope-anchored chromosome ends, that only a small fraction of the genome is nucleosomal, and that nucleosomes are re-established after replication without CAF-1.

## CONCLUSION

We have shown that at least some dinoflagellates retain all four core histone classes and candidate histone-modification pathways within an unconventional nuclear architecture. We propose a model in which a limited nucleosome population contributes predominantly to transcription-associated chromatin, alongside bulk DNA condensation involving non-histone proteins. The apparent reduction of canonical CAF-1 machinery is consistent with this model, but does not exclude histone functions in replication, repair or chromosome-end maintenance. Demonstrating assembled nucleosomes and mapping their occupancy against gene expression, DNA synthesis and nuclear position will be necessary to establish this proposed division of function.

## MATERIALS AND METHODS

### Nuclei isolation

*Oxyrrhis marina* and *Amphidinium carterae* were grown in F/2 seawater medium [74] supplemented with heat-killed bacteria (HKB) and F/2 seawater medium, respectively. Cells were harvested from 500 mL of culture by centrifugation at 3,000 × g for 10 min. Cells were incubated in either nuclear isolation buffer (NIB) adapted from Sikorskaite et al. [75] (10 mM HEPES-KOH, 10 mM NaCl, 10 mM KCl, 2.5 mM EDTA, 250 mM sucrose, 1 mM DTT, 0.5% NP-40), or from Sun et al. [76] (1 mM Tris, 1.5 mM CaCl_2_, 10 mM KCl, 0.1 mM DTT, 0.5% NP-40) for 10 min on ice with sonication (20% amplitude; 1 s on, 3 s off). Nuclei were then pelleted by centrifugation at maximum speed for 30 s. These steps were repeated with fresh NIB until the pellet was white. Samples were stored at −20 °C until proteomic processing.

For electron microscopy, *O. marina* nuclei were isolated as above with an additional Percoll/sucrose density-gradient purification step, in which the crude nuclear preparation was layered over 30% Percoll in NIB above a 2.5 M sucrose cushion. The gradient was centrifuged at 1,100 × g for 1 h, and the white layer that formed between the 30% Percoll/NIB and sucrose layers was collected. Nuclei were then washed in NIB without NP-40 and pelleted at 1,100 × g for 15 min.

### Electron Microscopy preparation

Isolated nuclei were fixed in NIB without NP-40 containing 2.5% glutaraldehyde overnight. For transmission electron microscopy (TEM), samples were post-fixed in 1% osmium tetroxide for 2 h and rinsed briefly in distilled water. Samples were stained en bloc with 0.25% uranyl acetate at 4 °C overnight. Samples were dehydrated through a graded acetone series (50%, 10 min; 70%, 2 × 10 min; 95%, 2 × 10 min; 100%, 2 × 10 min; dried 100%, 10 min). Samples were infiltrated with Epon–Araldite resin in dried acetone (3:1 acetone:resin for 3 h, then 1:3 overnight), followed by 100% resin for 6 h with one change after 3 h, embedded in fresh 100% resin and polymerised at 60 °C for 48 h. Thin sections (approximately 100 nm) were cut with a diamond knife on a Reichert–Jung Ultracut E ultramicrotome, collected on 300-mesh copper grids and stained with 2% aqueous uranyl acetate (10 min), rinsed in distilled water (2 × 5 min), stained with lead citrate (4 min), rinsed briefly in distilled water and air-dried. Sections were viewed with a JEOL JEM 1230 transmission electron microscope at 80 kV. Images were captured with a Hamamatsu ORCA-HR digital camera or with a Gatan 832 ORIUS SC1000 digital camera on an FEI Tecnai-12 microscope at 80 kV.

For scanning electron microscopy (SEM), glutaraldehyde-fixed nuclei were post-fixed in 1% osmium tetroxide for 2 h. The sample was then rinsed with distilled water and dehydrated through a graded ethanol series (50%, 10 min; 70% and 95%, 2 × 10 min each; 100%, 10 min; dried 100%, 2 × 10 min). Samples were critical-point dried in a Leica EM CPD300 automated critical point dryer, mounted on 13 mm aluminium stubs with double-sided carbon tabs or tape, and sputter-coated with gold/palladium in a Leica EM ACE600 high-vacuum coater at 30 mA. Samples were imaged with a ZEISS Sigma 300 field-emission SEM using a secondary electron (SE2) detector; the images shown were acquired at an accelerating voltage (EHT) of 5 kV and a working distance of 11.3 mm.

### Whole Cell Proteomic Sample Preparation

*Oxyrrhis marina* was grown in 500 mL of F/2 seawater medium supplemented with heat-killed bacteria (HKB). Once dense, the culture was split and grown for three days in F/2 medium, after which cells were pelleted by centrifugation at 3,000 × g for 10 min.

Protein was extracted from the cell pellet using 150 μL of a solution of 20 mM Tris-HCl pH 7.5, 150 mM KCl, 5 mM MgCl_2_, 0.5% (v/v) NP-40, 0.5% (w/v) sodium deoxycholate, and 0.5× complete protease inhibitor (Sigma). To support degradation of DNA/RNA, 10 μL of a nuclease solution (1 U/μL benzonase) was added to the lysate and incubated at 24 °C for 10 min. After nuclease digestion, 50 μL of a solution of 600 mM Tris-HCl pH 7.5, 8% (w/v) SDS, and 10 mM dithiothreitol (DTT) was added to the solution and incubated at 95 °C for 5 min in a heating block. After cooling to 24 °C, lysate was alkylated using 20 mM chloroacetamide (final concentration) and subsequently quenched with DTT. Protein concentration of the resulting lysate was measured using a BCA assay. Protein cleanup prior to digestion was carried out using an adapted version of the previously described SP3 protocol [77]. Briefly, 10 μL of a 50 mg/mL carboxylate-coated bead stock (Cytiva) was added to 40 μg of protein in 100 μL of solution followed by 400 μL of acetone. After a 5-min incubation at 24 °C, mixtures were centrifuged for 5 min at 5,000 × g. Resulting supernatants were discarded and pelleted material was resuspended in 800 μL of 80% ethanol and pelleted again as above. The ethanol supernatant was removed and 1 μg of trypsin/LysC in 100 μL of 4 M urea was added to each sample and incubated at 30 °C for 4 h with shaking at 800 rpm. Samples were then diluted to 1 M urea by addition of 300 μL of 100 mM Tris-HCl pH 7.5 and digested for an additional 16 h under the same conditions. After digestion, the supernatant was collected after centrifugation at 12,000 × g for 2 min and subsequently desalted using Strata-X SPE (Phenomenex, CAT# 8M-S100-4GA) micro-elution plates. SPE wells were initially conditioned with 100% (v/v) methanol (100 μL) followed by equilibration with 0.1% (v/v) trifluoroacetic acid (TFA) (100 μL). Loaded peptides were rinsed using 4% (v/v) methanol in 0.1% (v/v) formic acid (200 μL) and eluted using 60% (v/v) methanol in 0.1% (v/v) formic acid (100 μL). All SPE steps were carried out using a centrifuge with spinning at 250 × g for 3 min. Eluted peptides were evaporated to dryness in a SpeedVac centrifuge and reconstituted in a solution of 1% dimethylsulfoxide (v/v) in 1% (v/v) formic acid. Reconstituted peptide solutions were quantified on a QTRAP5500 MS using measured signal relative to a standard of known concentration.

### Data Independent Acquisition

Peptide samples were acquired using a data-independent acquisition (DIA) routine on an Orbitrap Fusion Lumos mass spectrometer (MS). Samples were introduced to the MS using a Dionex UltiMate 3000 liquid chromatography (LC) system equipped with a trapping-analytical column setup. For injection, peptides were initially trapped using 95% mobile phase A (0.1% formic acid) on a 100 μm inner diameter × 3 cm length column packed in-house with 1.9 μm Reprosil Pur C18 beads (Dr. Maisch, r119.aq.). Gradient elution of peptides was performed using a ramp of mobile phase B (80% v/v acetonitrile in 0.1% formic acid) on a 100 μm inner diameter × 25 cm length analytical column packed with 1.9 μm Reprosil Pur C18 beads. For each injection, 1 μg of peptides was separated with an 80-min linear LC gradient from 5% to 11% mobile phase B in 0.5 min, and to 34% B in 72.5 min at 400 nL/min, followed by rinsing and re-equilibration over the remaining 7 min. The analytical column outlet was coupled to a 20 μm inner diameter LOTUS electrospray tip (Fossil Ion Tech.). Each peptide sample was injected three individual times (80-min acquisition each), with each acquisition covering analysis of a separate mass range (injection 1 = 430–550, 2 = 550–690, 3 = 690–930 m/z). The complete duty cycle of the acquisition method consisted of two MS1 scans and two sets of windowed MS2 scans (order MS1 - MS2 - MS1 - MS2). For the first, low mass range injection, the initial MS1 scan covered a mass range of 425–555 m/z at a resolution of 60,000. The following set of DIA MS2 scans covered a precursor range of 430–550 m/z with an isolation window size of 4 m/z (0 m/z overlap) for a total of 30 scan events. Each scan used an HCD energy of 30 % and covered a defined mass range of 200–1,800 m/z at a resolution of 30,000. The next MS1 scan and following DIA MS2 used the same settings as the previous with the exception that the MS2 precursor mass range was set to 428–552 m/z to give a 2 m/z stagger with the previous scan windows. For the second, medium mass range injection, the scan settings were the same as for the low mass injection with the following changes: 1. MS1 scan range of 545–695 m/z; 2. DIA-MS2 precursor scan ranges of 550–690 m/z and 548–692 m/z; 3. Loop control number of spectra = 35. For the third, high mass range injection, the scan settings were the same as the others with the following changes: 1. MS1 scan range of 685–935 m/z; 2. DIA-MS2 precursor scan ranges of 690–930 m/z and 686–934 m/z; 3. DIA-MS2 isolation window of 8 m/z. All scan data were acquired in centroid mode.

### Proteomics for Nuclei Preparations

Three separate *O. marina* nuclear preparations (R1–R3) were isolated, pelleted and prepared for proteomics as described above for whole cells. DDA-MS analysis was carried out on an Orbitrap Lumos instrument interfaced to an UltiMate 3000 LC system as above. For this analysis, the gradient ramped from an initial starting condition of 5% mobile phase B (80% acetonitrile, 0.1% formic acid) to 34% B over 45 min, with sample pickup and column equilibration using an additional 15 min (60-min total run time, mobile phase A = 0.1% formic acid in water). Data acquisition on the Orbitrap Lumos utilized a standard data-dependent acquisition scheme. Specifically, the Orbitrap Lumos was globally set to use a positive ion spray voltage of 2,200 V, an ion transfer tube temperature of 275 °C, a default charge state of 2, advanced precursor determination activated, and an RF Lens setting of 45%. MS1 survey scans covered a mass range of 375–1,400 m/z at a resolution of 60,000 with an automatic gain control (AGC) target of 4e5 (100%) and the injection time set to ‘Auto’. Precursors for tandem MS/MS (MS2) analysis were selected using monoisotopic precursor selection (‘Peptide’ mode), charge state filtering (2–4z), dynamic exclusion (20 ppm low and high, 60 s duration, exclude isotopes = TRUE, exclude within cycle = TRUE), and an intensity threshold of 2.5e4. MS2 scans were carried out in the Orbitrap at a resolution of 30,000 using a higher-energy collision dissociation setting of 30%, an AGC target of 2e5 (400%), injection time set to ‘Auto’, and an isolation window setting of 1.6 m/z. The total allowable MS2 cycle time was set to 3 s. MS1 data were acquired in profile mode, and MS2 in centroid.

### Proteomic searches and quantification

Predicted proteomes from re-assembled MMETSP transcriptomes [35,36] were used as search databases for *Oxyrrhis marina* (MMETSP0044, MMETSP0451, MMETSP0468–MMETSP0471 and MMETSP1424–MMETSP1426) and *Amphidinium carterae* (MMETSP0258, MMETSP0259, MMETSP0398 and MMETSP0399). Sequences are referred to by assembly and gene or ORF identifier (e.g., MMETSP0469 g.127961); full identifiers are given in Tables S1–S4. After deduplication, initial searches of the three *O. marina* nuclear preparations were performed with basic FragPipe searches (MSFragger [78]), with label-free quantification by IonQuant [79]. The FragPipe DIA_SpecLib_Quant workflow [80] was used for the *O. marina* whole-cell sample and the *A. carterae* nuclear preparation, with quantification by IonQuant and DIA-NN [81]. Briefly, these searches allowed up to two missed cleavages, with methionine oxidation and protein N-terminal acetylation as variable modifications, and identifications were filtered at 1% FDR. The resulting identified proteins were then combined to create a reduced database for histone-specific (expanded) searches of each organism. These searches allowed up to four missed cleavages and five variable modifications per peptide, with the maximum peptide length lowered from 50 to 30 residues; acetylation, methylation and dimethylation were each allowed up to three times per peptide, and oxidation and phosphorylation up to twice. All samples were quantified with IonQuant, and DIA data additionally with DIA-NN.

To estimate the contribution of core histones to the proteome, razor-peptide IonQuant intensities of proteins assigned to each histone family were summed per class and divided by the summed razor intensity of all proteins in the run. DVNP and HLP fractions were calculated in the same way, and enrichment was expressed relative to the whole-cell sample [Tables S2, S3]. Histone peptides identified in the dinoflagellate datasets were compared with the human proteome, and peptides with 100% identity to any human protein were flagged as potential contaminants; peptides not shared with human histones are marked with triangles in Figures S2 and S3.

### Protein annotation and subcellular localisation

Proteins identified in the *O. marina* whole-cell sample and R1 were annotated with InterProScan [37] (E ≤ 10^−5^); proteins without an InterProScan match were classed as unannotated. Subcellular localisation was predicted with DeepLoc [38]. For Figure 1B, razor intensities were summed by predicted localisation (nucleus, cytoplasm or other) and annotation status [Table S1].

### Comparative survey of chromatin-associated proteins

Homologues of core and linker histones, DVNP, HLP, histone chaperones and assembly factors, and histone-modifying enzymes [Figure 1A] were sought in predicted proteomes from UniProt [82] (*Perkinsus marinus*, *Vitrella brassicaformis*, *Homo sapiens*, *Tetrahymena thermophila*, *Ectocarpus siliculosus*, *Plasmodium falciparum*, *Giardia lamblia* and *Trypanosoma brucei*), EukProt [83] (*Amoebophrya* sp. and *Hematodinium* sp.) and re-assembled MMETSP transcriptomes [35,36] (*O. marina*, *A. carterae*, *L. polyedra* and *K. veneficum*), using homology- and profile hidden Markov model-based searches [search tools, versions, query sequences and thresholds to be added]. Absences were recorded separately for genome- and transcriptome-derived datasets. Accessions and sequences are listed in Table S4.

### Histone curation and phylogenetic analysis

Candidate core histones (H2A, H2B, H3 including CENP-A, and H4) were retrieved from proteomes predicted with TransDecoder [84] from the re-assembled MMETSP transcriptomes [35,36] with BLASTP 2.12.0 [85] (BLOSUM62, e-value ≤ 10^−5^), using human H2A type 1-B/E (P04908), H2B type 1-J (P06899), H3.1 (P68431) and H4 (P62805) as queries. Sequences of *Hematodinium* sp. SG-2012 and *Chromera velia* were taken from EukProt [83]. Histones detected by proteomics in *O. marina* and *A. carterae* (25 sequences) were identified with BLASTP (e-value ≤ 10^−5^) and confirmed with InterProScan 5.73-104.0 [37].

Each histone family was processed separately. Terminal stop codons were removed, sequences were trimmed to the first methionine when it lay within the first 25 residues, and sequences shorter than 90 or longer than 300 amino acids were discarded; UniProt references were not trimmed or filtered. Redundancy was reduced within each species only, pooling all MMETSP strains of a species, so that conserved histones shared between species were retained. In each species, the sequence most similar to the human canonical histone (H2A type 1, P0C0S8; H2B type 1-J, P06899; H3.1, P68431; H4, P62805), measured as identical positions in a global BLOSUM62 alignment in Biopython 1.83 [86], was always kept. Sequences at least 90% identical to it were removed with cd-hit-2d, and the rest were clustered at 90% identity with CD-HIT 4.8.1 [87,88]. Proteomics-identified sequences were then added. Reference core histones and their variants (including H2A.Z, H2AX, macroH2A, H3.3 and CENP-A/CENH3) were obtained from UniProt [82] on 24 October 2026 and were clustered per organism at 90% identity with the same anchor procedure.

Strains were screened for contamination using COX1 transcripts, identified with BLASTP (BLOSUM62, e-value ≤ 10^−5^) using UniProt COX1 sequences as queries. The 386 COX1 contigs (85 strains) were searched with BLASTN 2.12.0 (e-value ≤ 10^−10^) against a local database of 84,820 dinoflagellate and non-dinoflagellate COX1 and mitochondrial genome sequences from NCBI GenBank (14 October 2026), and the three best hits per contig were classified by NCBI Taxonomy [89]. After manual review, sequences from strains with discordant COX1 hits were relabelled in the trees and excluded from the root-to-tip analysis.

For the full histone phylogeny, the four family datasets were aligned with MAFFT 7.505 L-INS-i [90] and trimmed with trimAl 1.5.1 (-gappyout) [91]. A maximum-likelihood tree was inferred with IQ-TREE 3.0.1 [92] under Q.insect+F+R9 [93,94], selected by the Bayesian information criterion in ModelFinder [95]. Branch support was estimated with 1,000 SH-like approximate likelihood ratio test replicates [96] and 1,000 ultrafast bootstrap replicates [97]. An H3/CENP-A tree was inferred in the same way under Q.insect+G4 [98].

Root-to-tip distances (patristic, in substitutions per site) were measured on the full histone tree, rooted at the midpoint of the branch separating H2A and H2B from H3 and H4. Dinoflagellates were compared with representative metazoa and with the proteomics-identified histones of *O. marina*; possible contaminants and tips outside their family clade were excluded. Distributions were plotted as Gaussian kernel density estimates with Matplotlib 3.6.3 [99].

### Gene density along contigs

Gene density towards contig ends was determined from the genome annotations of two of the most contiguous dinoflagellate assemblies available, *Cladocopium goreaui* SCF055 (GCA_947184155.2) and *Durusdinium trenchii* CCMP2556 (GCA_963970005.1). The longest 20% of contigs were retained for analysis (*Cladocopium goreaui*: 1,369 contigs ≥255 kb; *Durusdinium trenchii*: 5,828 contigs ≥13 kb; *Polarella glacialis* was excluded owing to its lower contiguity). For each gene, the relative position of its midpoint along the contig (gene midpoint/contig length; 0 and 1 represent the two ends) was calculated. Gene density (genes per Mb) was calculated in equal-width bins of relative position. Gene distributions were tested with a Kolmogorov–Smirnov test and a paired Wilcoxon signed-rank test [Figure S7].

## DATA AVAILABILITY

Protein identifications, quantifications and candidate modifications are provided in Tables S1–S5. Proteomic files and all other scripts, results or methods can be accessed through Figshare.

## ACKNOWLEDGEMENTS

We acknowledge Mary Ann Trevors from the Electron Microscopy Core Facility at Dalhousie University for expert assistance with sample preparation and technical guidance in scanning and transmission electron microscopy. We extend our thanks to Ping Li (Scientific Imaging Suite, Dalhousie University) for operational assistance with the FEI Tecnai-12 TEM, and to Christopher Hughes for constant and insightful proteomic support provided through the Biological Mass Spectrometry Core Facility at Dalhousie University; and Susana Breglia for maintaining dinoflagellate cultures and advice regarding sample preparation.

## SUPPLEMENTARY MATERIAL

Figure S1. BUSCO completeness (v6.1.0, alveolate_odb12.2) of the dinoflagellate MMETSP proteomes.

Figure S2. High-scoring histone peptide-spectrum matches from initial and expanded FragPipe searches of *O. marina* whole cells and nuclear preparations. Annotated MS/MS spectra are shown for H4, H3, H2A and H2B peptides (b ions, green; y ions, orange). Modified peptides are marked with stars and peptides that are not identical to any human histone sequence with triangles.

Figure S3. High-scoring histone peptide-spectrum matches from initial and expanded FragPipe searches of the *A. carterae* nuclear preparation. Symbols as in Figure S2.

Figure S4. High-scoring DVNP and HLP peptide-spectrum matches from initial and expanded FragPipe searches of *O. marina* whole cells and nuclear preparations and the *A. carterae* nuclear preparation. Methylated and dimethylated peptides are labelled.

Figure S5. Electron microscopy of isolated *O. marina* nuclei. Transmission electron micrographs of isolated nuclei show condensed, chromosome-like profiles throughout a granular nucleoplasm (top left and top right), with a higher-magnification view (middle right) and a lower-magnification field of the preparation (bottom left, upper panel). Scanning electron micrographs show a constricted, paired body (bottom left, lower panel) and an ovoid body with pore-like surface openings (arrows; bottom right).

Figure S6. Scanning electron micrographs of isolated *O. marina* nuclear preparations. (A) Bodies with relatively smooth or folded surfaces, including lobed and constricted forms. (B) Bodies with granular surfaces, including paired and clustered forms. (C) Higher-magnification views of surface texture and pore-like openings. Images were acquired at 5 kV; scale bars and magnifications are given in each image data bar.

Figure S7. Gene density along the longest contigs of the *Cladocopium goreaui* and *Durusdinium trenchii* genome assemblies. Points show gene density (genes per Mb) in equal-width bins of relative position along contigs (0 and 1, contig ends), and lines show smoothed fits. Kolmogorov– Smirnov (KS) and paired Wilcoxon test results are given in each panel.

Figure S8. Alignment of a partial *O. marina* DVNP with viral DVNP homologues (UniProt accessions shown); residues with candidate modifications in *O. marina* are red.

Table S1. DeepLoc-predicted localisation, razor intensity and InterPro annotation of proteins identified in the *O. marina* whole-cell sample and R1, with summed intensities by predicted localisation.

Table S2. *O. marina* histone and DVNP identifications from initial and expanded histone searches of nuclear preparations R1–R3 and the whole-cell sample: sequence coverage, peptides, razor peptides, PSMs, IonQuant (total, unique and razor) and DIA-NN intensities, histone and DVNP fractions of summed intensity, fold enrichment and candidate modifications.

Table S3. *A. carterae* histone, DVNP and HLP identifications from the nuclear preparation, with the same fields as Table S2.

Table S4. Presence (1) or absence (0) of DNA-packaging, nucleosome-assembly and histone-modification components across the 14 taxa in Figure 1A, with accessions and sequences.

Table S5. Nucleoporin homologues identified in the *O. marina* proteome, grouped by nuclear pore complex subcomplex.

