## Supplemental_Figures_S1-7 for "Proteomics reveals conserved features of histone biology in dinoflagellates"

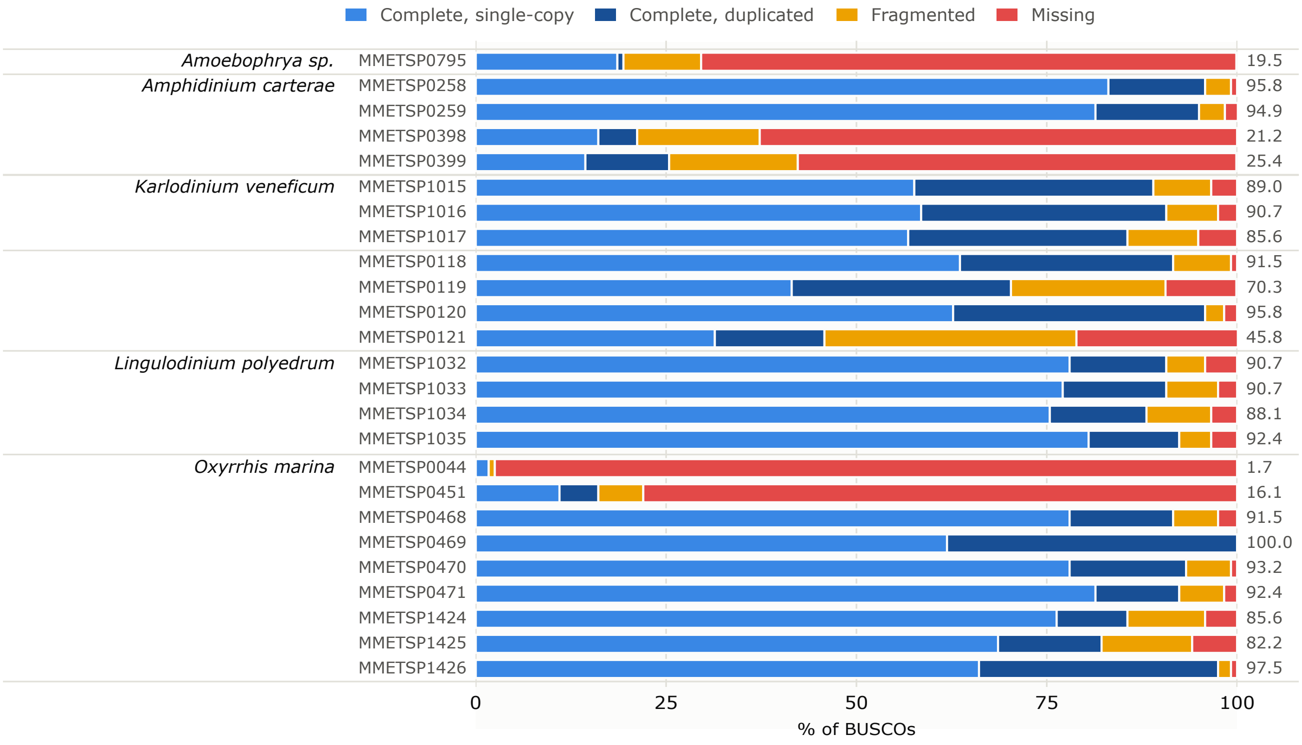

Complete, single-copy
Complete, duplicated
Fragmented
Missing
Amoebophrya sp.
MMETSP0795
19.5
Amphidinium carterae
MMETSP0258
95.8
MMETSP0259
94.9
MMETSP0398
21.2
MMETSP0399
25.4
Karlodinium veneficum
MMETSP1015
89.0
MMETSP1016
90.7
MMETSP1017
85.6
MMETSP0118
91.5
MMETSP0119
70.3
MMETSP0120
95.8
MMETSP0121
45.8
Lingulodinium polyedrum
MMETSP1032
90.7
MMETSP1033
90.7
MMETSP1034
88.1
MMETSP1035
92.4
Oxyrrhis marina
MMETSP0044
1.7
MMETSP0451
16.1
MMETSP0468
91.5
MMETSP0469
100.0
MMETSP0470
93.2
MMETSP0471
92.4
MMETSP1424
85.6
MMETSP1425
82.2
MMETSP1426
97.5
0
25
50
75
100
% of BUSCOs

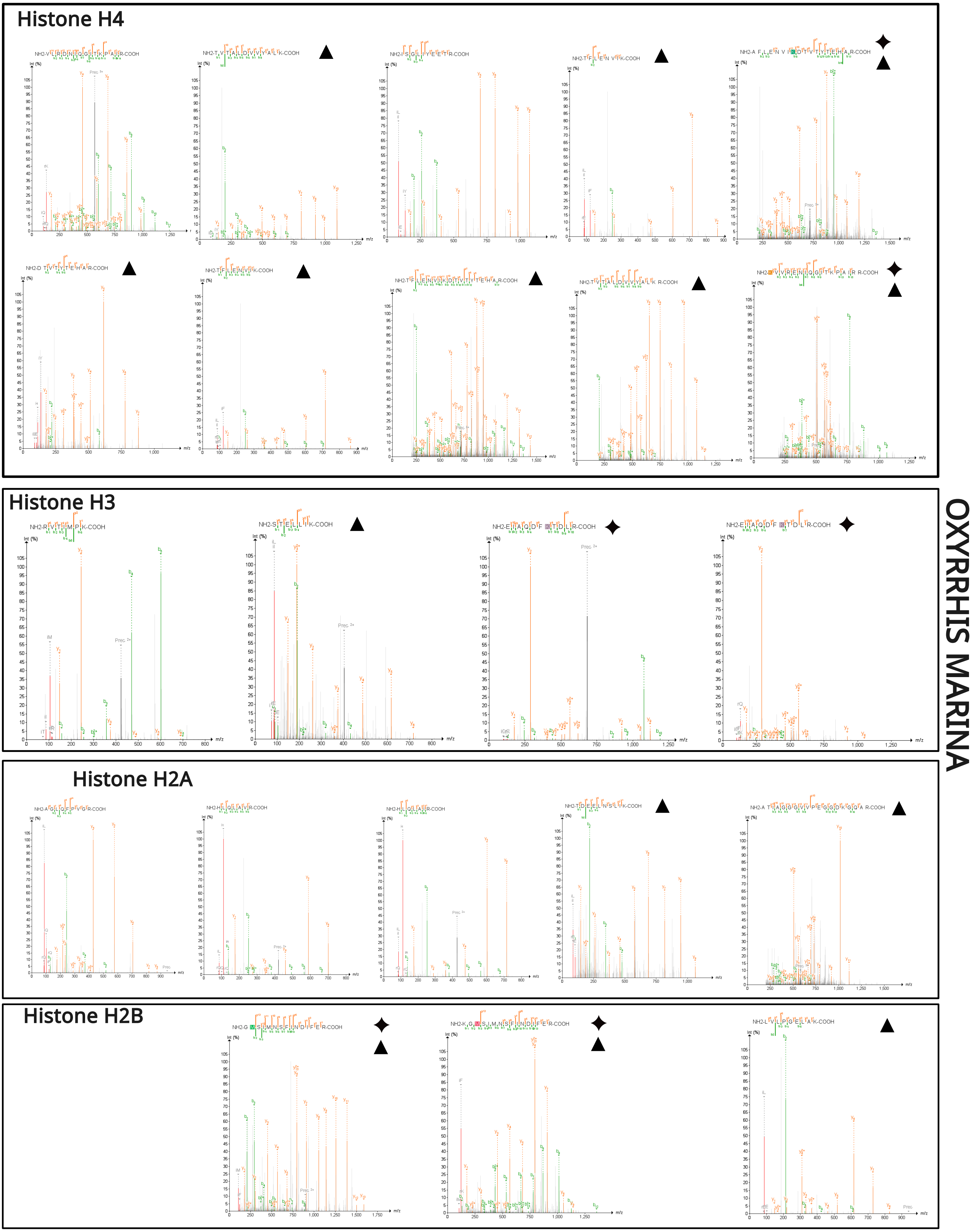
Histone H4
Histone H3
Histone H2A
Histone H2B
OXYRRHIS MARINA

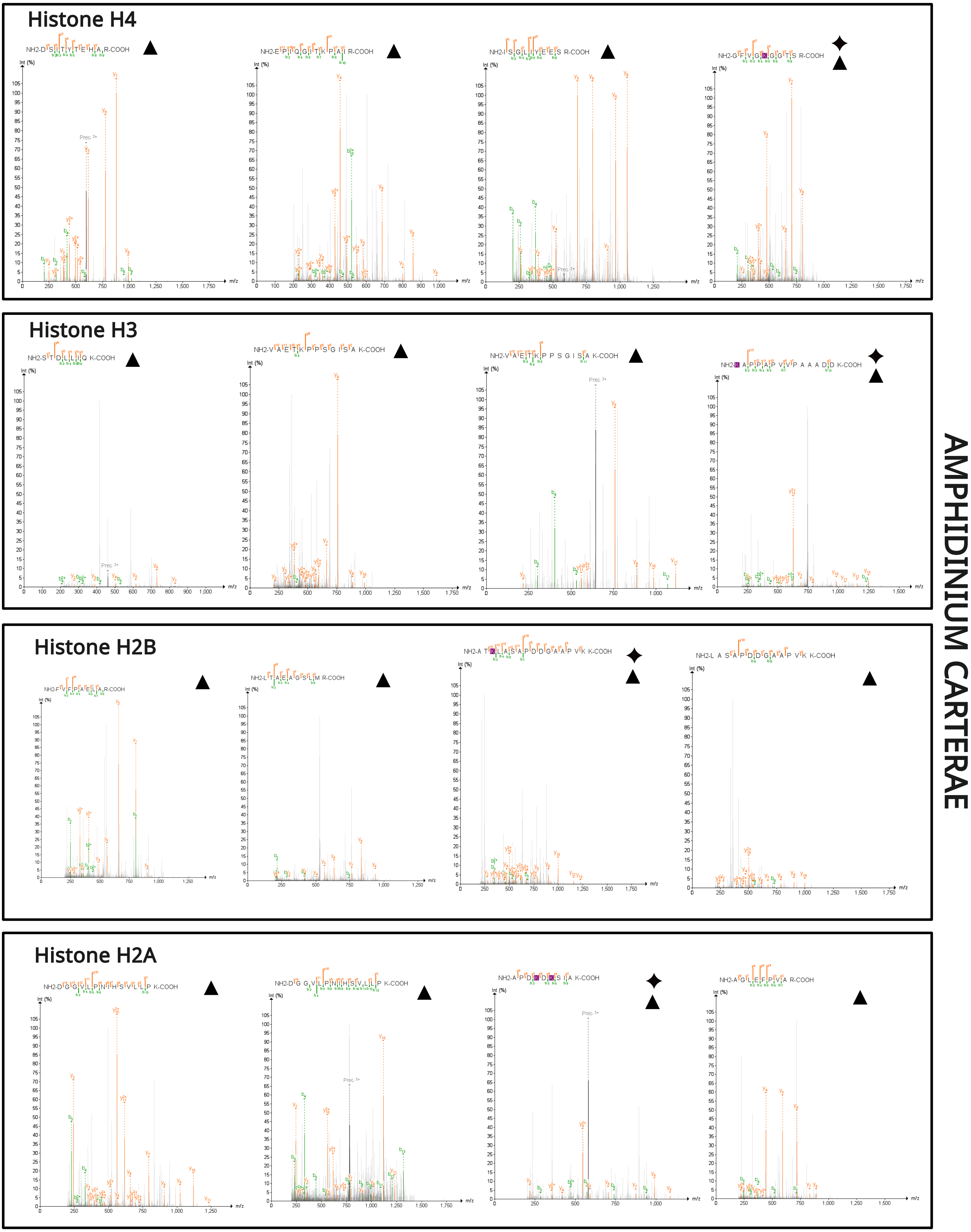
Histone H4
Histone H3
Histone H2B
Histone H2A
AMPHIDINIUM CARTERAE
Int (%)
m/z

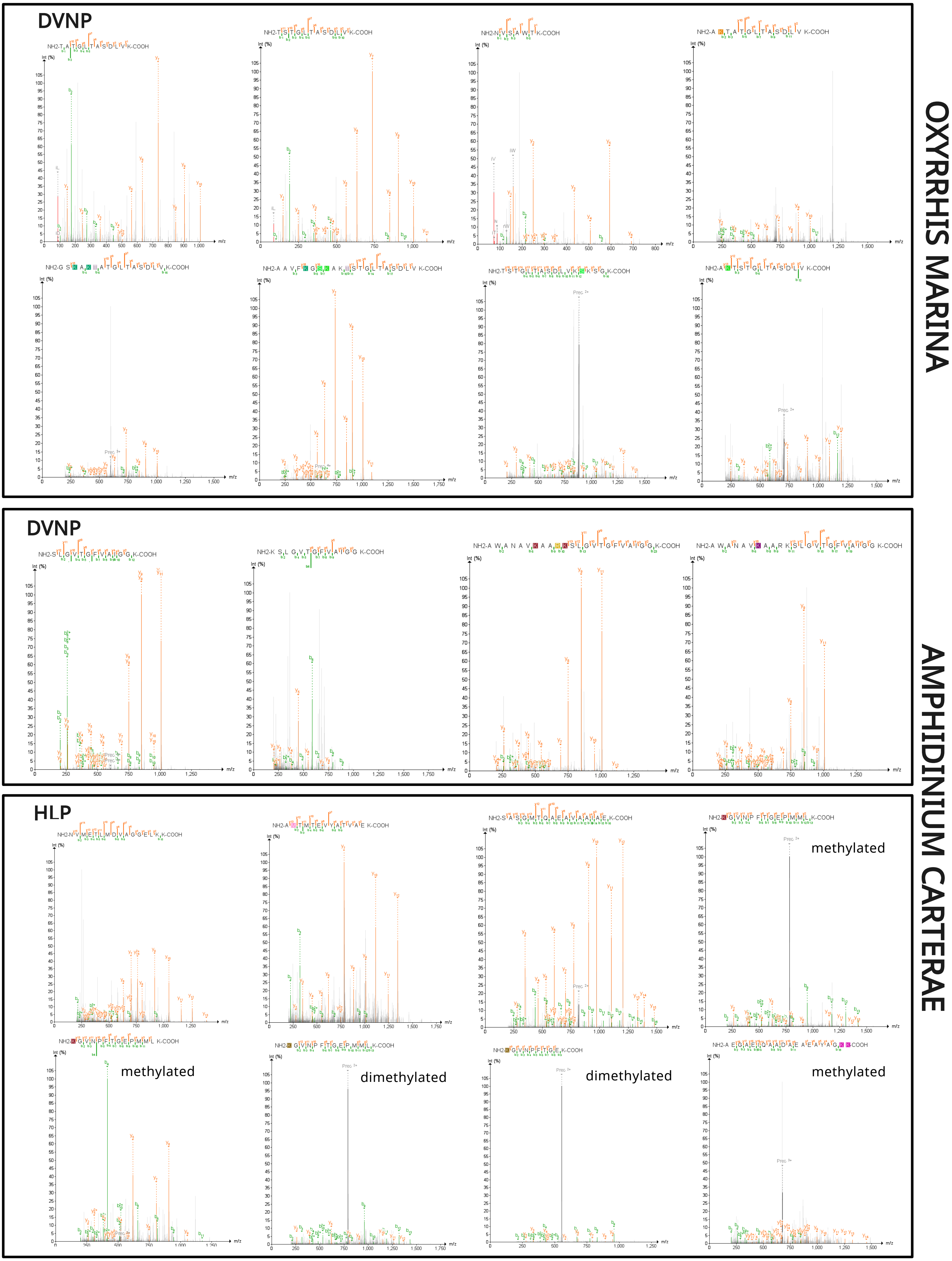
DVNP
OXYRRHIS MARINA
DVNP
AMPHIDINIUM CARTERAE
HLP
methylated
methylated
dimethylated
dimethylated
methylated

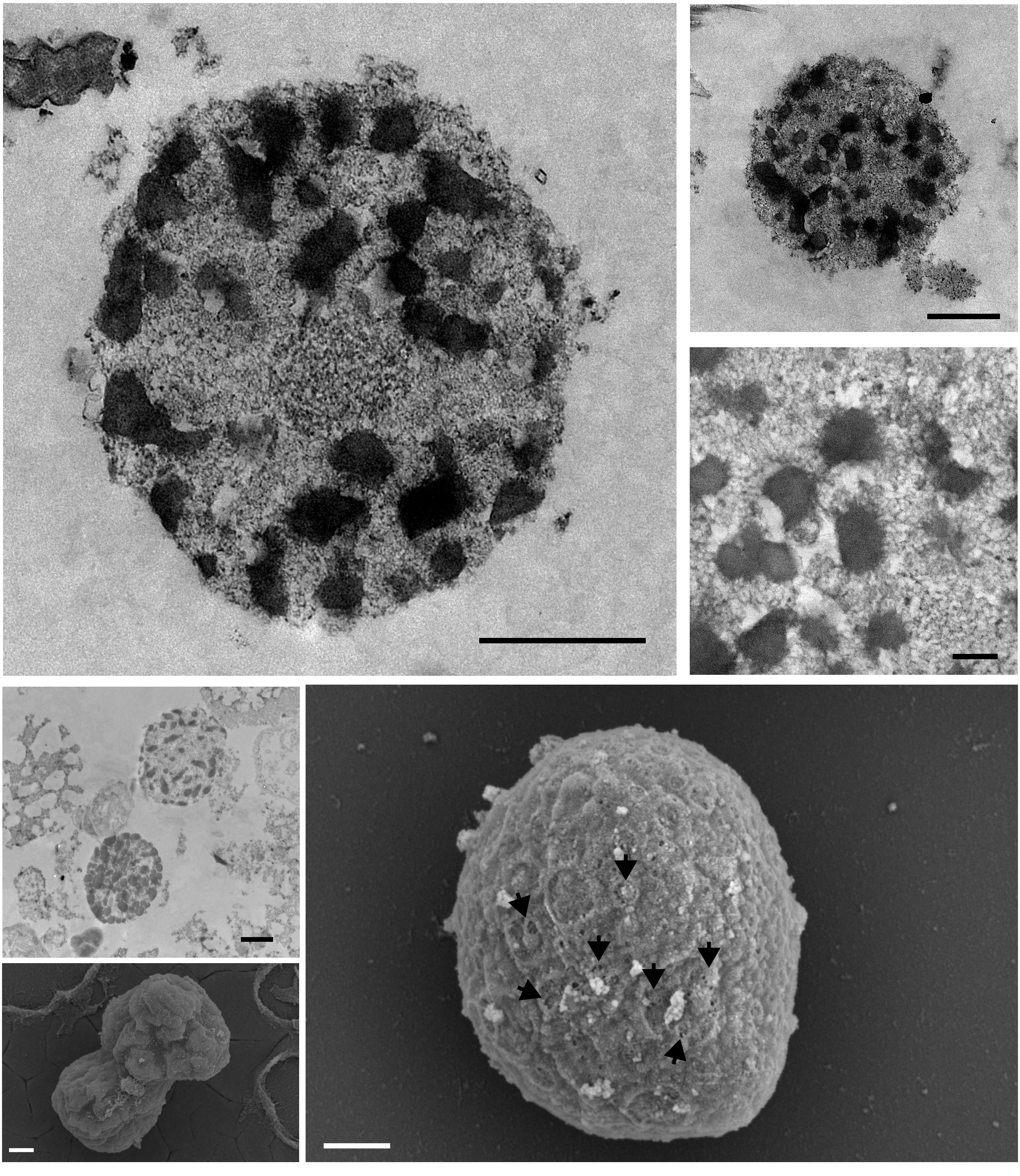

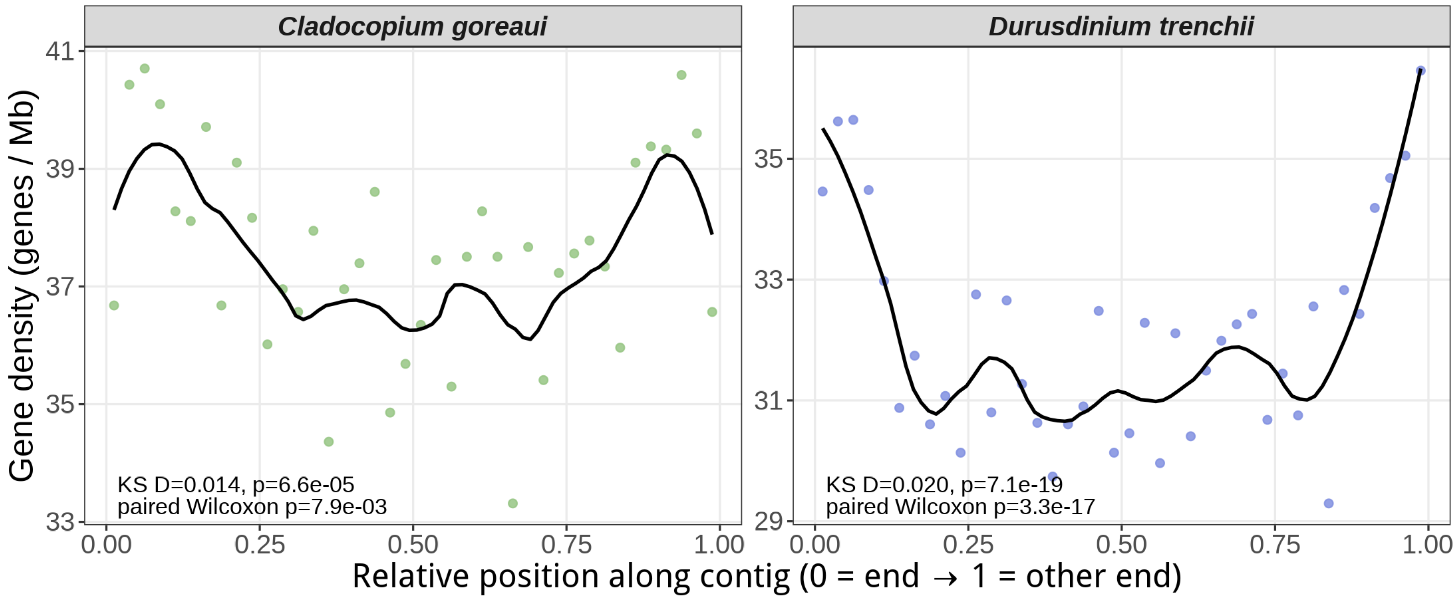
Cladocopium goreaui
Durusdinium trenchii
Gene density (genes / Mb)
41
39
37
35
33
35
33
31
29
KS D=0.014, p=6.6e-05
paired Wilcoxon p=7.9e-03
KS D=0.020, p=7.1e-19
paired Wilcoxon p=3.3e-17
0.00
0.25
0.50
0.75
1.00
0.00
0.25
0.50
0.75
1.00
Relative position along contig (0 = end → 1 = other end)

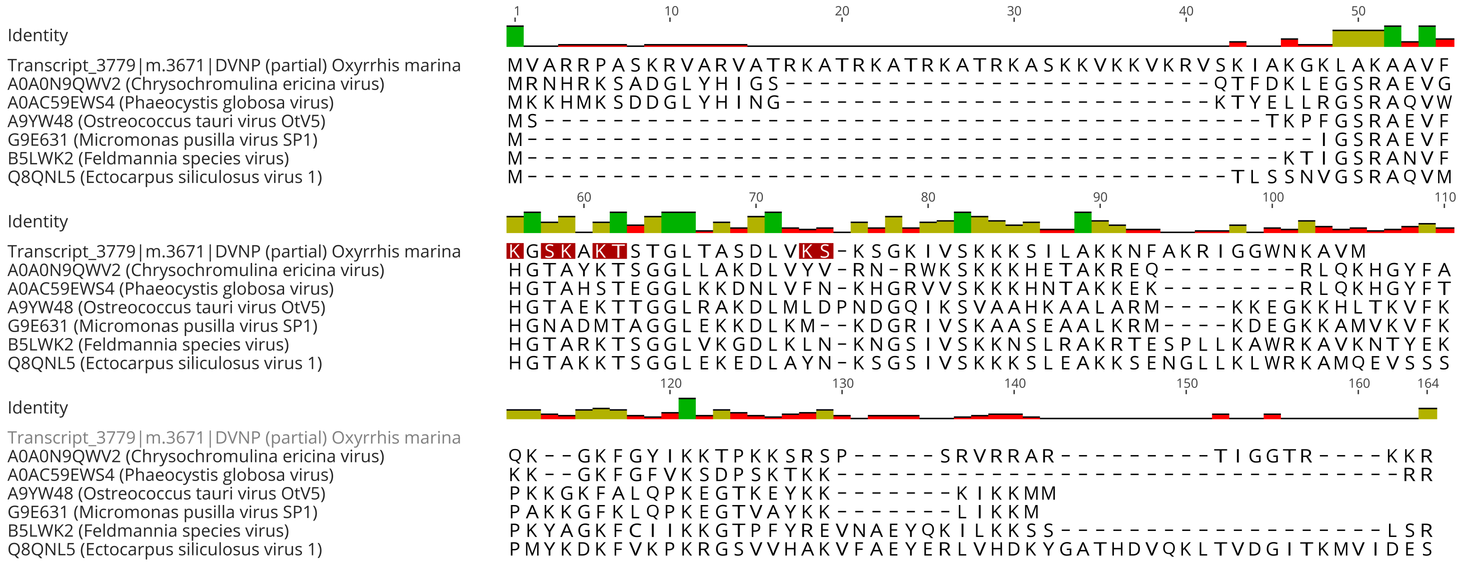

1 10 20 30 40 50
Identity
Transcript_3779|m.3671|DVNP (partial) Oxyrrhis marina MVARRPASKRVARVATRKATRKATRKATRKASKKVKKVKRVSKIAKGKLAKAAVF
A0A0N9QWV2 (Chrysochromulina ericina virus) MRNHRKSADGLYHIGS-------------------------QTFDKLEGSRAEVG
A0AC59EWS4 (Phaeocystis globosa virus) MKKHMKSDDGLYHING-------------------------KTYELLRGSRAQVW
A9YW48 (Ostreococcus tauri virus OtV5) MS------------------------------------------TKPFGSRAEVF
G9E631 (Micromonas pusilla virus SP1) M----------------------------------------------IGSRAEVF
B5LWK2 (Feldmannia species virus) M--------------------------------------------KTIGSRANVF
Q8QNL5 (Ectocarpus siliculosus virus 1) M-----------------------------------------TLSSNVGSRAQVM
60 70 80 90 100 110
Identity
Transcript_3779|m.3671|DVNP (partial) Oxyrrhis marina KGSKAKTSTGLTASDLVKS-KSGKIVSKKKSILAKKNFAKRIGGWNKAVM
A0A0N9QWV2 (Chrysochromulina ericina virus) HGTAYKTSGGLLAKDLVYV-RN-RWKSKKKHETAKREQ--------RLQKHGYFA
A0AC59EWS4 (Phaeocystis globosa virus) HGTAHSTEGGLKKDNLVFN-KHGRVVSKKKHNTAKKEK--------RLQKHGYFT
A9YW48 (Ostreococcus tauri virus OtV5) HGTAEKTTGGLRAKDLMLDPNDGQIKSVAAHKAALARM----KKEGKKHLTKVFK
G9E631 (Micromonas pusilla virus SP1) HGNADMTAGGLEKKDLKM--KDGRIVSKAASEAALKRM----KDEGKKAMVKVFK
B5LWK2 (Feldmannia species virus) HGTARKTSGGLVKGDLKLN-KNGSIVSKKNSLRAKRTESPLLKAWRKAVKNTYEK
Q8QNL5 (Ectocarpus siliculosus virus 1) HGTAKKTSGGLEKEDLAYN-KSGSIVSKKKSLEAKKSENGLLKLWRKAMQEVSSS
120 130 140 150 160 164
Identity
Transcript_3779|m.3671|DVNP (partial) Oxyrrhis marina
A0A0N9QWV2 (Chrysochromulina ericina virus) QK--GKFGYIKKTPKKSRSP-----SRVRRAR---------TIGGTR----KKR
A0AC59EWS4 (Phaeocystis globosa virus) KK--GKFGFVKSDPSKTKK---------------------------------RR
A9YW48 (Ostreococcus tauri virus OtV5) PKKGKFALQPKEGTKEYKK-------KIKKMM
G9E631 (Micromonas pusilla virus SP1) PAKKGFKLQPKEGTVAYKK-------LIKKM
B5LWK2 (Feldmannia species virus) PKYAGKFCIIKKGTPFYREVNAEYQKILKKSS-------------------LSR
Q8QNL5 (Ectocarpus siliculosus virus 1) PMYKDKFVKPKRGSVVHAKVFAEYERLVHDKYGATHDVQKLTVDGITKMVIDES
